# *Dictyostelium discoideum* cells sense their local density and retain nutrients when the cells are about to overgrow their food source

**DOI:** 10.1101/2022.04.08.487657

**Authors:** Ramesh Rijal, Sara A. Kirolos, Ryan J. Rahman, Richard H. Gomer

**Author notes:** Address correspondence to: Richard H. Gomer, Department of Biology, Texas A&M University, ILSB 301 Old Main Drive, College Station, TX 77843-3474.

## Abstract

*Dictyostelium discoideum* is a unicellular eukaryote that eats bacteria, and eventually overgrows the bacteria. *D. discoideum* cells accumulate extracellular polyphosphate (polyP), and the polyP concentration increases as the local cell density increases. At high cell densities, the correspondingly high extracellular polyP concentrations allow cells to sense that they are about to overgrow their food supply and starve, causing the *D. discoideum* cells to inhibt their proliferation. In this report, we show that high extracellular polyP inhibits exocytosis of undigested or partially digested nutrients. PolyP decreases cell membrane fluidity and plasma membrane recycling, and this requires the G protein-coupled polyP receptor GrlD, the polyphosphate kinase Ppk1, and the inositol hexakisphosphate kinase I6kA. PolyP did not affect random cell motility, cell speed, or F-actin levels. PolyP decreased membrane saturated fatty acids and altered lipid and protein contents in detergent-insoluble lipid microdomains. Together, these data suggest that *D. discoideum* cells use polyP as a signal to sense their local cell density and reduce cell membrane fluidity and membrane recycling, perhaps as a mechanism to retain ingested food when the cells are about to starve.

## Introduction

Some animals use seasonal cues to anticipate starvation during winter, and store food as body fat for hibernation (Florant and Healy, 2012). A similar process occurs in the unicellular eukaryote *Dictyostelium discoideum*, which uses polyphosphate (linear chains of phosphate groups) as a secreted signal to monitor the local cell density, and when there is a high density of cells that will soon overgrow the local food supply, the cells stop proliferating but continue to grow (accumulate mass and protein) in anticipation of starvation (Soll et al., 1976; Yarger et al., 1974). PolyP inhibits proliferation by inhibiting cytokinesis, but has relatively little effect on growth (Suess and Gomer, 2016). In yeast, there exist mutations that inhibit proliferation but do not inhibit growth (Johnston et al., 1977), and conversely some embryos during early development have cell proliferation without growth (Johnston et al., 1977; Nasmyth, 1996; Su and O’Farrell, 1998). However, much remains to be understood about how cells separately regulate growth and proliferation.

*D. discoideum* is a soil dwelling amoeba that shares common ancestors with plants and animals (Baldauf and Doolittle, 1997). During growth, the cells feed on bacteria and divide. When starved, *D. discoideum* cells aggregate to form fruiting bodies consisting of amass of spore cells held above the ground by a column of stalk cells. Dispersal of spores to a moist environment causes the spores to hatch and germinate into amoeba (Kessin, 2001; Loomis, 2014; Schaap, 2011). For nutrient reserves, *D. discoideum* cells store lipid in the form of lipid droplets (Kornke and Maniak, 2017), and glycogen, which is broken down during development (Garrod and Ashworth, 1972; Harris and Rutherford, 1976).

Polyphosphate (polyP) is present in many cell types (Rao et al., 2009). In prokaryotes, polyP is synthesized from ATP by polyphosphate kinase 1 (Ppk1) (Ahn and Kornberg, 1990; Brown and Kornberg, 2008), and is involved in stress responses, virulence, quorum sensing, and biofilm formation (Rao et al., 2009). PolyP is also found in archaea and eukaryotes (Lander et al., 2013; Orell et al., 2012; Ruiz et al., 2004). In eukaryotic cells, polyP is both secreted and present in the cytoplasm, vacuoles, nucleus, mitochondria, and plasma membranes (Beauvoit et al., 1989; Lichko et al., 2006; Livermore et al., 2016; Offenbacher and Kline, 1984; Reusch, 1992; Suess and Gomer, 2016; Urech et al., 1978; Verhoef et al., 2017).

PolyP is present in *D. discoideum* acidocalcisomes (electron-dense acidic calcium storage organelles involved in intracellular pH homeostasis and osmoregulation (Docampo et al., 2005; Marchesini et al., 2002), contractile vacuoles, mitochondria, nuclei, and the cytoplasm (Gezelius, 1974; Marchesini et al., 2002; Satre et al., 1986). *D. discoideum* also accumulate extracellular polyP (Suess and Gomer, 2016). At high cell densities, the concomitant high extracellular concentrations of polyP inhibit *D. discoideum* proliferation and induce aggregation, the first stage of development (Suess and Gomer, 2016; Suess et al., 2017). PolyP thus appears to be a signal that cells use to sense their local cell density to anticipate starvation. *D. discoideum* cells use the G protein-coupled receptor GrlD to bind and sense polyP (Suess and Gomer, 2016; Suess et al., 2019). In proliferating *D. discoideum* cells, extracellular polyP accumulation is regulated by polyphosphate kinase 1 (Ppk1) and inositol hexakisphosphate kinase A (I6kA) (Suess and Gomer, 2016). Wild type *D. discoideum* cells accumulate intracellular polyP during starvation (Livermore et al., 2016), and cells lacking Ppk1 are multinucleate and form less spores than wild type cells (Livermore et al., 2016), suggesting that polyP affects both growth and development.

In this report, we find that extracellular polyP reduces cell membrane fluidity and recycling, and alters the composition of lipid microdomains. Possibly as a result, polyP inhibits exocytosis of ingested food particles in *D. discoideum* cells which appears to cause cells to retain food when they are about to overgrow their food supply and starve. This mechanism thus allows polyP to simultaneously promote growth and inhibit proliferation.

## Results

### PolyP inhibits exocytosis in *D. discoideum*

*D. discoideum* cells accumulate extracellular polyP as their cell density increases, and the extracellular polyP concentration (≥ 470 μg/ml), characteristic of a high cell density, inhibits *D. discoideum* cell proliferation by inhibiting cytokinesis without affecting the growth of the cells (Suess and Gomer, 2016). This then causes cells to be larger and have more protein/cell at stationary phase than in mid-log phase (Soll et al., 1976), and thus to have more stored nutrients in anticipation of starvation. Another possible way for cells to store nutrients is to prevent digestion of endocytosed nutrients, and/or prevent excretion (by exocytosis) of partially digested nutrients. *D. discoideum* cells can endocytose dextran, a non degradable fluid phase marker (Hacker et al., 1997; Klein and Satre, 1986; Rivero and Maniak, 2006), and the dextran is then exocytosed after typically 15-30 minutes (Klein and Satre, 1986). To determine if polyP promotes retention of ingested material, cells were exposed to a 30 minute pulse of tetramethylrhodamine isothiocynate (TRITC)-dextran and were then washed free of extracellular TRITC-dextran. PolyP concentrations greater than or equal to 470 μg/ml increased retention of ingested TRITC-dextran after 30 minutes of exocytosis (Figures 1A-B and S1A-C). Other sources and preparations of polyP including 2-kDa filtered polyP, short chain polyP (< 60 -mer) medium chain (~100-mer), and 60-mer polyP also increased the retention of ingested TRITC-dextran (Figures 1C and S1D). *D. discoideum* cells require GrlD to bind polyP (Suess et al., 2019), and require Ppk1 and i6kA to accumulate extracellular polyP (Suess and Gomer, 2016). To test if polyP uses a signal transduction pathway to induce retention of TRITC-dextran, cells lacking GrlD (*grlD*^−^), Ppk1 (*ppk1*^−^), I6kA (*i6kA*^−^), and *i6kA*-null cells overexpressing I6kA (*i6kA^−^/i6kA*) were assayed. Compared to wild-type (WT) cells, 470 and 705 μg/ml polyP did not increase the retention of TRITC-dextran in *grlD*^−^, *ppk1*^−^, and *i6kA*^−^ cells, but the effect of polyP was partially rescued in *i6kA^−^/i6kA* cells (Figures S1E and F), suggesting that polyP uses a signal transduction pathway involving GrlD, Ppk1, and I6kA to inhibit exocytosis. To determine if the increased retention of TRITC-dextran in WT and *i6kA^−^/i6kA* cells was due to increased macropinocytosis of TRITC-dextran, WT, *grlD*^−^, *ppk1*^−^, *i6kA^−^* and *i6kA^−^/i6kA* cells were incubated with TRITC-dextran, and the levels of ingested TRITC-dextran were measured. PolyP at 470 and 705 μg/ml decreased macropinocytosis in WT but did not significantly alter macropinocytosis in *grlD*^−^, *ppk1*^−^, and *i6kA*^−^ cells (Figures S1G and H). Overexpressing I6kA in *i6kA^−^/i6kA* cells rescued the phenotype at 470 but not 705 μg/ml (Figures S1G and H), suggesting that polyP uses a signal transduction pathway involving GrlD and Ppk1, and possibly I6kA to inhibit macropinocytosis. Together, the data suggest that polyP decreases ingestion, but increases retention of ingested food. At 470 μg/ml polyP, the ingestion decreases by 29% and the retention increases by 32%, while at 705 μg/ml polyP the ingestion decreases by 14% and the retention increases by 21%, which would cause the net mass of the cell to increase over time, which has been previously observed (Suess and Gomer, 2016; Yarger et al., 1974)

**Figure 1.**
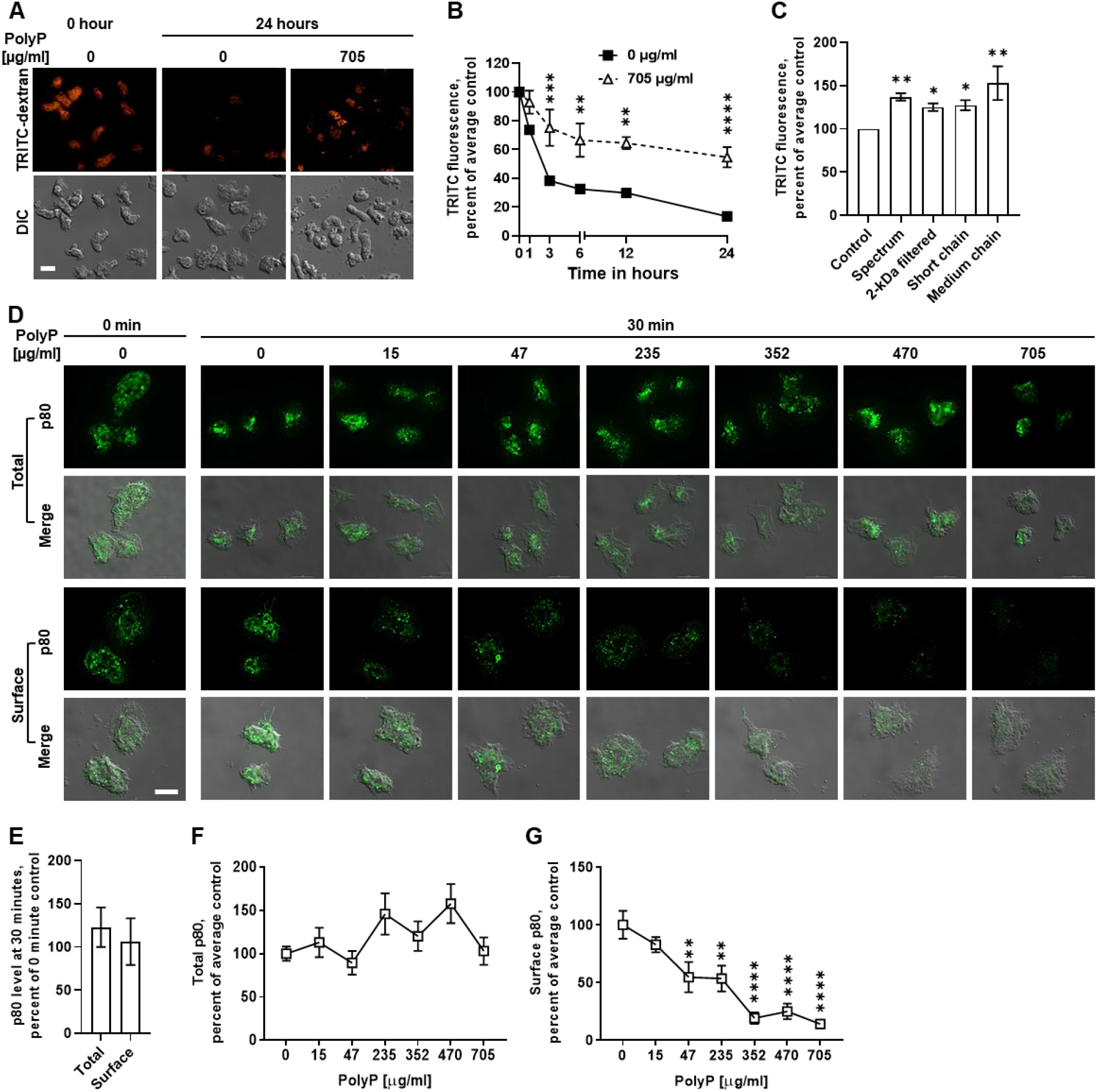
PolyP inhibits exocytosis in *D. discoideum*. **A)** WT *D. discoideum* cells were incubated with TRITC-dextran in the absence (0) or presence of 705 μg/ ml polyP, the uningested TRITC-dextran was removed by washing, and cells were imaged at 0 and 24 hours. DIC indicates differential interference contrast. Bar is 10 μm. Images are representative of 3 independent experiments. **B)** Quantification of TRITC-dextran fluorescence from **A**. The average of no polyP (0) was considered 100%. **C)** TRITC-dextran fluorescence per cell in the absence or presence of 705 μg/ ml of the indicated polyP at 30 minutes. **D)** Immunofluorescence images of cells stained for p80 (green), with fixed and permeabilized cells in the top two rows, and fixed but not permeabilized cells in the bottom two rows. Bar is 10 μm. Images are representative of 3 independent experiments. **E)** Fluorescence intensity from permeabilized cells (total p80) or non-permeabilized cells (surface p80) at time 30 minutes in **D** was normalized to fluorescence intensity from permeabilized cells (total p80) or non-permeabilized cells (surface p80) at time 0 minute, repectively. **F)** Quantification of fluorescence intensity from permeabilized cells (total p80) in **D**. **G)** Quantification of fluorescence intensity from non-permeabilized cells (cell surface p80) in **D**. In F and G, the fluorescence intensity of p80 with no polyP (0) was considered 100%. All values in B, C, E, F, and G are mean ± SEM of 3 independent experiments. * indicates p < 0.05, ** p < 0.01, *** p < 0.001, **** p < 0.0001 comparing polyP to no polyP at each time in B, compared to control in C, and compared to 0 polyP in G (Šídák’s multiple comparisons test (B), Holm-Šídák’s multiple comparisons test (C and D)).

Extracellular polyP at concentrations ranging from 5 to 15 μg/ml, which correspond to the concentrations of polyP in medium cell density cultures, inhibit the killing of ingested *Escherichia coli* (*E. coli*) in *D. discoideum* cells without significantly affecting the ingestion of *E. coli* or fluorescently labeled heat-killed yeast (zymosan) bioparticles (Rijal et al., 2020). As previously observed for 15 μg/ml polyP, 47, 470, and 705 μg/ml polyP inhibited the killing of ingested *E. coli* by *D. discoideum* cells at 24 hours after ingestion (Figure S1I). 705 μg/ml polyP also reduced the number of ingested zymosan bioparticles (Figure S1J).

Together, these data suggest that the extracellular polyP concentrations characteristic of high cell densities somewhat inhibit endocytosis and phagocytosis, more strongly inhibit exocytosis, and also inhibit the killing of ingested *E. coli*. The net result is that high extracellular polyP concentrations cause a retention, and to some extent preservation, of ingested nutrients.

During exocytosis, late endosomes fuse with the plasma membrane and form transient exocytic membrane microdomains (Charette and Cosson, 2006). These microdomains are enriched with the transmembrane protein p80, and thus p80 at the plasma membrane can be used to monitor exocytosis in *D. discoideum* (Charette and Cosson, 2006). To determine if polyP causes the retention of TRITC-dextran by inhibiting exocytosis, we examined total and cell-surface p80. The levels of p80 did not change when cells were incubated for 30 minutes in the absence of polyP (Figures 1D and E). Exposure of cells to polyP for 30 minutes did not significantly affect levels of total p80, but 47 μg/ml and higher polyP decreased surface p80 levels (Figure 1D-F), suggesting that polyP inhibits the surface localization of p80 in *D. discoideum* cells, possibly due to a reduction in exocytosis.

### PolyP increases the retention of internalized membranes

*D. discoideum* cells grow and proliferate in liquid medium and uptake nutrients by pinocytosis (Vines and King, 2019), which requires the continuous internalization of the plasma membrane. However, cells maintain their total surface area by membrane recycling, which involves exocytic fusion of the internalized plasma membrane back to the plasma membrane (Thilo and Vogel, 1980). *D. discoideum* cells replace the entire plasma membrane every ~45 minutes by internalization and exocytosis (Thilo and Vogel, 1980). To determine if polyP inhibition of exocytosis inhibits cell membrane recycling, we incubated WT cells with polyP for 30 minutes, the plasma membranes were stained with CellMask Green, a fluorescent lipid analogue, and images were taken. PolyP did not significantly affect the total cell staining (Figure 2A and B), whereas a 7 to 8 minute exposure of cell to 587 or 705 μg/ml polyP increased the accumulation of the fluorescent lipid in the interior of cells (Figures 2A and C). This suggests that polyP increases the retention of internalized cell membranes.

**Figure 2.**
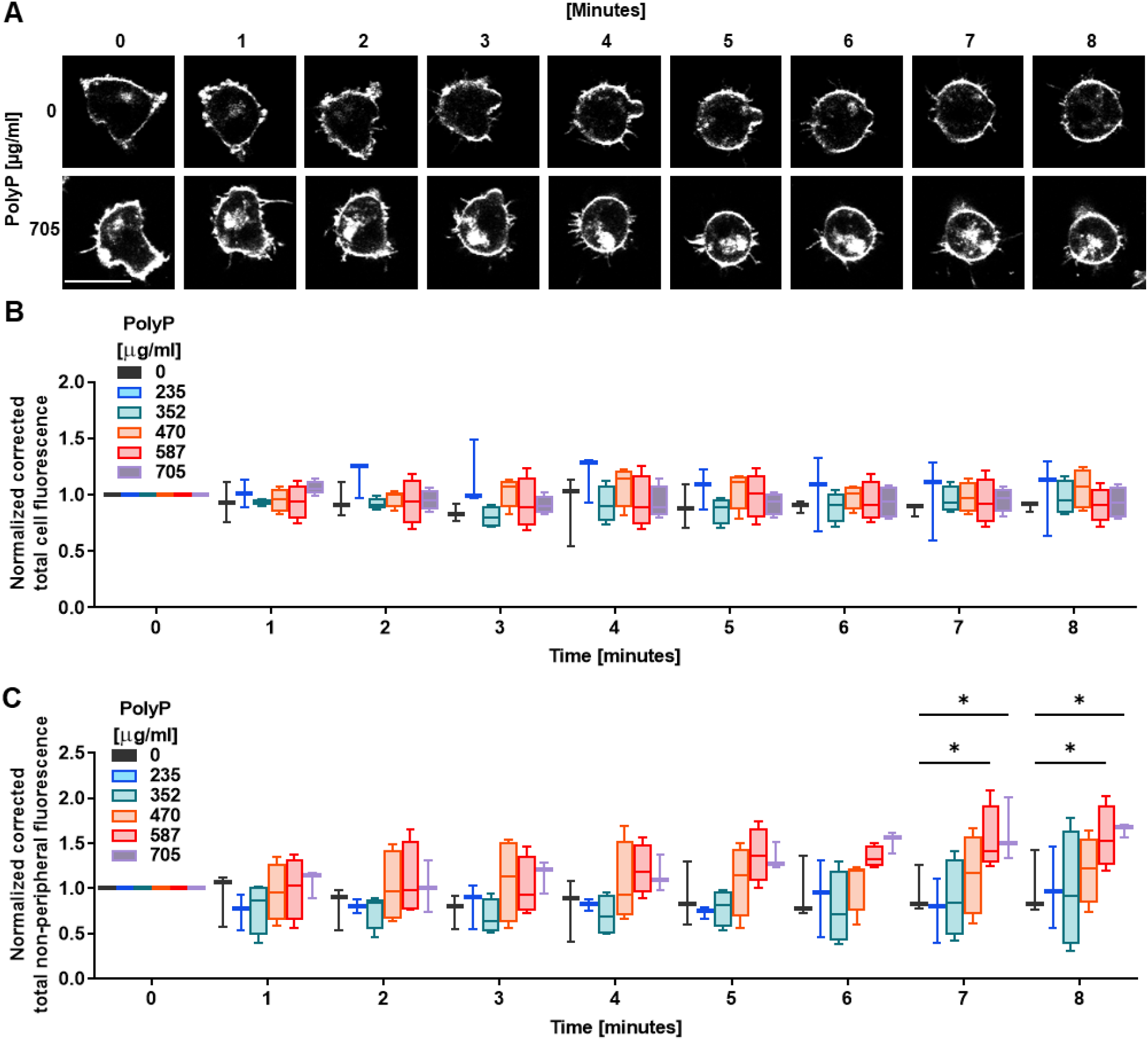
PolyP increases retention of internalized membranes in the cells. **A-C)** WT *D. discoideum* cells were incubated with the indicated concentration of polyP, stained with membrane dye CellMask green (gray), and total (A and B) and non-peripheral (A and C) fluorescence intensities were measured at the indicated times. Bar is 10 μm (A). All values are mean ± SEM of 3 independent experiments (B and C). * p < 0.05 (Dunnett’s multiple comparisons test).

### PolyP requires GrlD, Ppk1, and I6kA to reduce the cell membrane fluidity of WT *D. discoideum* cells

Physical properties of cell membranes, such as membrane fluidity, are critical determinants of efficient endocytosis and exocytosis in mammalian cells (Ben-Dov and Korenstein, 2013; Ge et al., 2010). To investigate if polyP alters cell membrane fluidity as a mechanism of inhibiting endocytosis and exocytosis, *D. discoideum* cells were stained with the membrane dye CellMask Green, photobleached with a high intensity laser beam, and the recovery of the fluorescence within the photobleached area in cells was measured. Compared to control with no polyP, 705 μg/ml Spectrum, 2-kDa filtered, short chain, and medium chain polyP increased the half-life of recovery of fluorescence after photobleaching by ~ two-fold in WT *D. discoideum* cells (Figures 3A-C and Movies 1 and 2), and decreased the diffusion coeffient (Figure 3D). The area beneath the curve in Figure 3B describes the mobile membrane fraction. Compared to the control, all polyP types reduced the membrane mobile fraction (Figure 3E). Spectrum polyP caused similar effects at concentrations ≥ 470 μg/ml (Figure S2A). Together, these data suggest that high cell density polyP (≥ 470 μg/ml) reduces the cell membrane fluidity of WT *D. discoideum* cells, and that the effect of polyP does not depend on chain length, source, or purity of the polyP.

**Figure 3.**
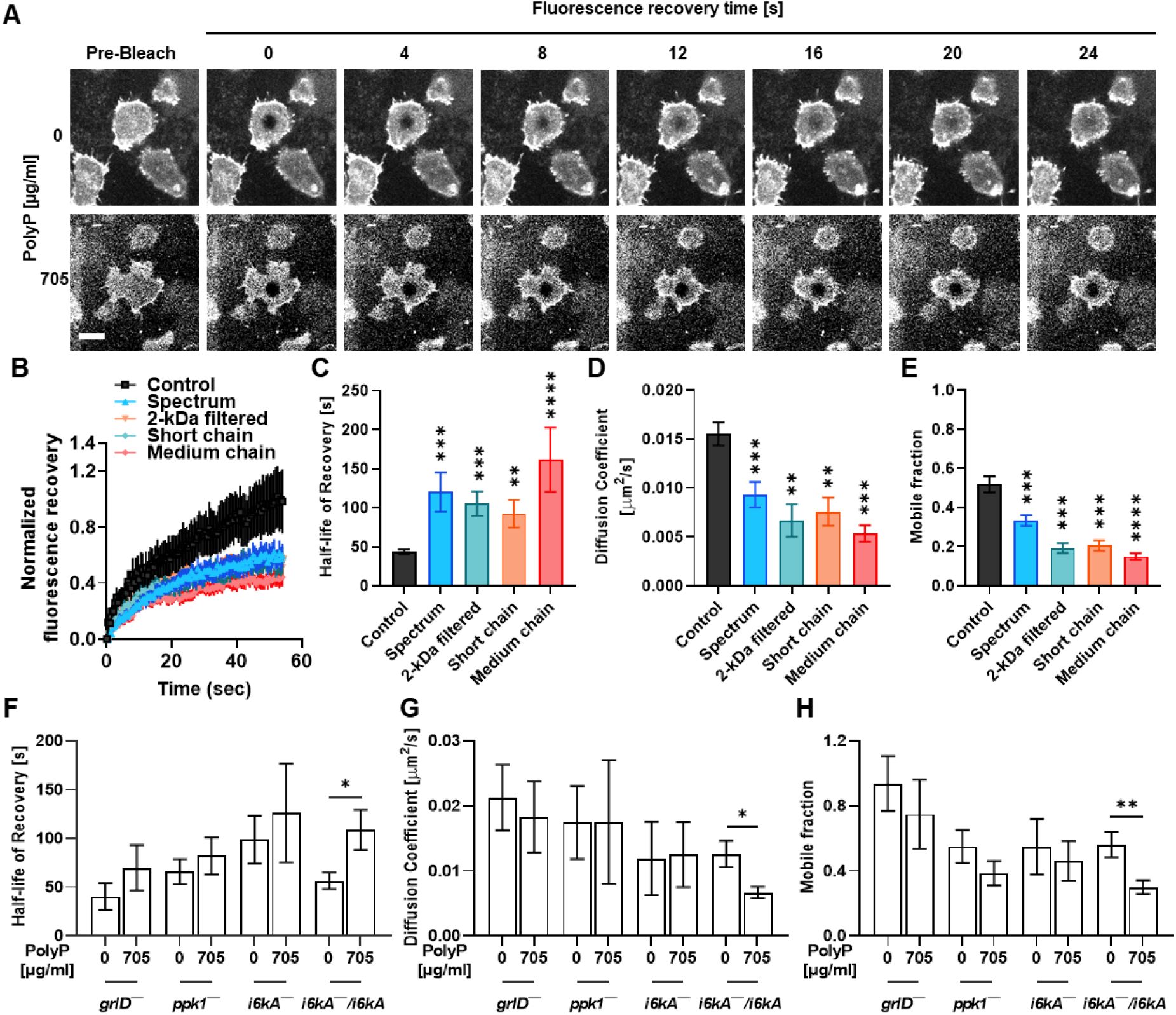
PolyP reduces the cell membrane fluidity of WT *D. discoideum* cells, and this effect of polyP requires GrlD, Ppk1, and I6kA. **(A)** WT *D. discoideum* cells were incubated with the indicated concentration of polyP for 30 minutes, stained with CellMask green (gray), photobleached using a 488 nm laser, and fluorescence recovery in the photobleached area was monitored over time. Bar is 10 μm. **(B)** Cells were incubated in the absence (control) or presence of 705 μg/ml of the indicated polyP for 30 minutes, and fluorescence recovery as in **A** was measured. Fluorescence intensity at 0 seconds in the bleached spot after photobleaching was considered 0. **(C-E)** Half-life of recovery, diffusion coefficient, and mobile fraction were generated from the data in **B**. **(F-H)** Half-life of recovery, diffusion coefficient, and mobile fraction were calculated for *grlD*^−^, *ppk1*^−^, *i6kA*^−^, and *i6kA^−^/i6kA* as in **B-E**. All values are mean ± SEM of at least 3 independent experiments. * p < 0.05, ** p < 0.01, *** p < 0.001, **** p < 0.0001 (Dunn’s multiple comparisons test (C-E) and Mann Whitney test (F-H)).

The conditioned medium (CM) from cells at high cell densities contains polyP (Suess and Gomer, 2016). To determine if CM can mimic the effect of exogenous polyP on membrane fluidity, CM was collected from high cell density (> 15 x 10^6^ cells/ml) WT cell cultures, and dilutions of CM were added to mid-log phase cells. Similar to exogenous polyP, ≥ 60% CM increased the half-life of recovery, and decreased the diffusion coefficient and mobile fraction in WT cells (Figure S2B), indicating that a factor present in the CM from cultures at high cell densities reduces cell membrane fluidity.

To determine if polyP inhibits membrane fluidity using a signal transduction pathway, we tested the effect of polyP on cell membrane fluidity of *grlD*^−^, *ppk1*^−^, *i6kA*^−^ and *i6kA^−^/i6kA* cells. Compared to the control with no polyP, 705 μg/ml polyP did not significantly change the half-life of recovery, diffusion coefficient, and mobile fraction of *grlD*^−^, *ppk1*^−^, and *i6kA*^−^ cells (Figure 3F-H). However, polyP increased the half-life of recovery, and decreased the diffusion coefficient and mobile membrane fraction of *i6kA^−^/i6kA* cells (Figure 3F-H). Interestingly, compared to WT cells, *i6kA*^−^ cells showed an increase in the half-life of recovery and a decrease in the diffusion coefficient, indicating inherently decreased membrane fluidity (Figures S2C and D). Similar to polyP, 100% CM did not significantly affect the cell membrane fluidity of *grlD*^−^, *ppk1*^−^, and *i6kA*^−^ cells (Figure S2C-E), but increased the half-life of recovery and decreased the diffusion coefficient and mobile fraction of *i6kA /i6kA* cells (Figures S2D and E), suggesting that exogenous polyP or high cell density CM uses a signal transduction pathway involving GrlD, Ppk1, and I6kA to reduce membrane fluidity in *D. discoideum* cells.

### PolyP does not alter random cell motility, speed, and cytoskeletal actin, but reduces directionality and increases the formation of filopodia

To determine if polyP-mediated reduced cell membrane fluidity affects cell motility, WT cells in the absence or presence of 705 μg/ml polyP were tracked for 30 minutes, and the accumulated distance (total distance travelled along its path), speed (accumulated distance divided by time), and directionality (the straight-line distance between the start and end of a cell’s movement divided by the accumulated distance) of cells were measured. Although polyP did not significantly affect accumulated distance and speed (Figures 4A-C), polyP reduced the directionality (Figure 4D). However, cells incubated with 100% CM showed reduced accumulated distance, speed, and directionality (Figure S3A-D), which may be due to the effect of unknown factors in the CM.

**Figure 4.**
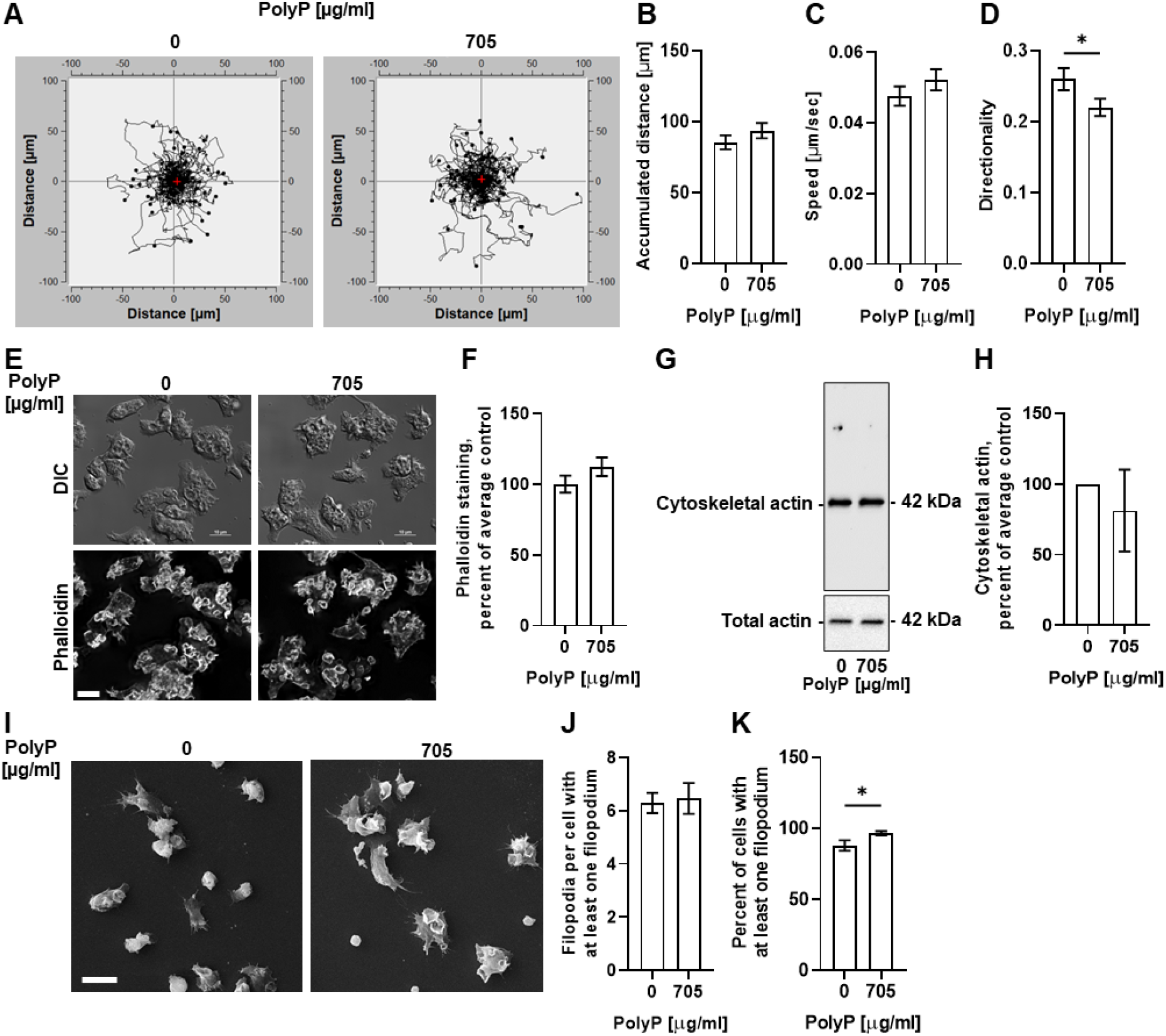
PolyP does not alter random cell motility, speed, and cytoskeletal actin, but reduces directionality, and increases the percent of the cells with filopods. **A)** WT cells in the absence (0) or presence of (705 μg/ml) polyP were filmed for 30 minutes; 30 cells per experiment were tracked, and tracks were graphed. Red plus sign indicates the center of mass after 30 minutes. The tracks are a compilation of three independent experiments with at least 30 tracks per experiment. **B-D)** Quantifications of the effect of polyP from A on cell displacement (accumulated distance), speed, and cell persistence (directionality) over 30 minutes. **E)** Differentital interference contrast (DIC) and fluorescence images of wild-type *D. discoideum* cells cultured for 30 minutes in the presence of the indicated concentration of polyP, and then stained with phalloidin (gray) for Factin. Images are representative of 3 independent experiments. Bar is 10 μm. **F)** Quantification of mean fluorescence intensity of phalloidin in **E**. The average of no polyP (0) was considered 100%. **G)** WT cells were incubated with the indicated concentration of polyP for 30 minutes, and Western blots of whole cell lysates or detergent-insoluble cytoskeletons were stained with anti-actin antibodies. Molecular masses in kDa are at right. Blots are representative of three independent experiments. **H)** Densitometry was used to estimate levels of polymerized actin. Polymerized actin densitometry was normalized to the total actin. The average of no polyP (0) was considered 100%. **I)** Scanning electron micrographs of WT cells cultured the indicated concentration of polyP for 30 minutes. Images are representative of 3 independent experiments. Bar is 10 μm. **J and K)** The effect of polyP on the number of filopodia per cell projecting at least one filopodium (J) and the percent of cells with at least one filopodium (K). All values are mean ± SEM from 3 independent experiments. * in D and K indicate p < 0.05 (Mann Whitney test).

To determine if extracellular polyP affects cytoskeletal actin to reduce cell membrane fluidity, WT cells were incubated with or without 705 μg/ml polyP for 30 minutes. The cells were then fixed and stained with phalloidin to determine the level of cytoskeletal actin. PolyP did not discernably change the distribution of, or significantly change the levels of, phalloidin staining (Figure 4E and F). The polyP also did not alter the level of cytoskeletal actin as assayed by western blots of crude cytoskeletons stained for actin (Figure 4G and H). Unlike 705 μg/ml polyP, 100% CM reduced phalloidin staining and cytoskeletal actin levels (Figure S3E-H), which as above may be due to the effect of unknown factors in the CM. Together, these data suggest that polyP does not alter cytoskeletal actin while reducing cell membrane fluidity.

Filipodia are actin rich protrusions involved in sensing the environment and cell anchorage on a surface (Heid et al., 2005; Wood and Martin, 2002). To determine if extracellular polyP affects filopodia while reducing cell membrane fluidity, we incubated WT cells with 0 or 705 μg/ml polyP for 30 minutes, and examined filopodia using scanning electron microscopy of fixed cells. PolyP slightly increased the percent of cells having at least one filopodium without affecting the number of filopodia per cell in the cells that did have filopodia (Figures 4I-K). Together, these results indicate that in addition to reducing cell membrane fluidity, polyP slightly decreases directionality of cell movement, causes an increase in the number of cells with actin-rich filopodia, but does not significantly affect cell speed or cytoskeletal actin.

### PolyP alters the lipid composition in lipid microdomains

Cell membrane specific lipid microdomains, also called lipid rafts, have been observed in different organisms ranging from bacteria to humans (Henderson and Block, 2014; Kaiser et al., 2009; Lopez, 2015). The organization and composition of lipids in the lipid microdomains are necessary to maintain cell membrane fluidity and integrity, which influences many biological events such as intracellular signaling (Ikonen, 2001; Simons and Toomre, 2000), cell adhesion and migration (Gomez-Mouton et al., 2001), and pathogenesis and immunity (Hartlova et al., 2010; Van Laethem and Leo, 2002). To determine if polyP changes the fatty acid and lipid composition of the lipid microdomains, cells were incubated with or without 705 μg/ml polyP for 30 minutes, lysed in 1% Triton X-100 to extract the Triton X-100 insoluble lipid microdomains, and the microdomains were analyzed by mass spectrometry (Metcalffe and Wang, 1981). In the control- and polyP-treated Triton X-100 insoluble lipid microdomains, triglycerides with fatty acids of varying chain lengths were the most abundant lipids (Table 1). Compared to control, polyP decreased the abundance of the saturated fatty acids hexadecanoic acid (also called palmitic acid) and stearic acid (Figure 5). Long chain fatty acids containing diglycerides, triglycerides, phosphatidyl glycerol, phosphatidyl ethanolamine, and phosphatidyl inositol were present only in the control (Table 2), whereas lipids containing short chain fatty acids such as diglycerides, triglycerides, ceramides, and phosphatidyl inositols were present only in polyP treated Triton X-100 insoluble lipid microdomains (Table 2). However, polyP increased level of the triglyceride (4:0_8:0_10:1), and decreased levels of the triglycerides (4:0_8:0_11:1) and (4:0_12:0_12:2) in the Triton X-100 insoluble fraction (Table 1). Together, these data indicate that polyP alters fatty acid and lipid compositions of the Triton X-100 insoluble lipid microdomains, which might contribute to a reduction in cell membrane fluidity.

**Figure 5.**
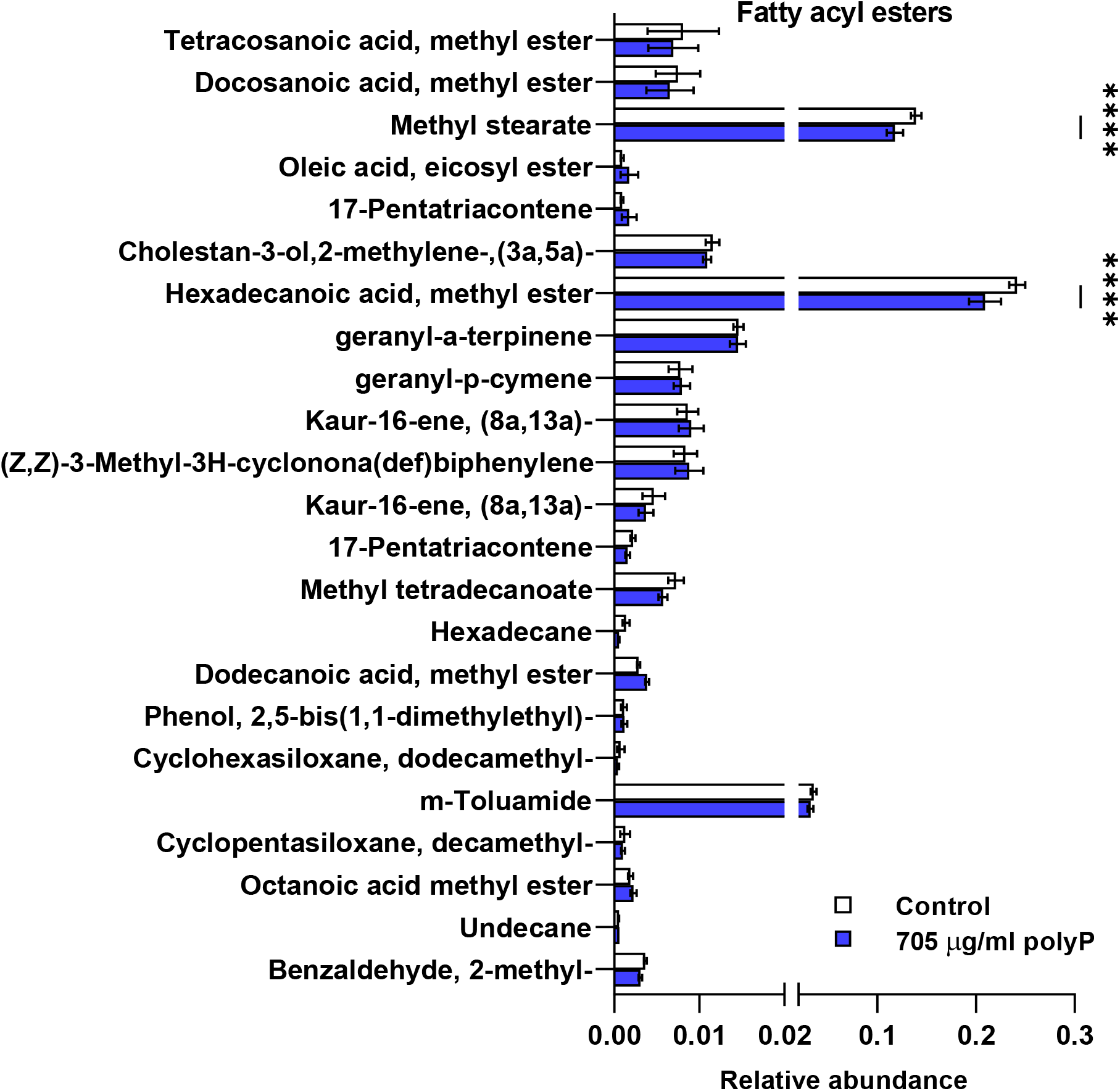
PolyP reduces some saturated fatty acids in detergent-insoluble lipid microdomains. Triton X-100 insoluble membranes were extracted from WT *D. discoideum* cells in the absence (control) or presence of (705 μg/ml) polyP and subjected to gas chromatography followed by mass spectrometry, and the abundances of fatty acyl esters relative to decanoic acid standard (not shown) were determined. All values are mean ± SEM of 3 independent experiments. **** indicates p < 0.0001 (Bonferroni’s multiple comparisons test).

**Table 1:**
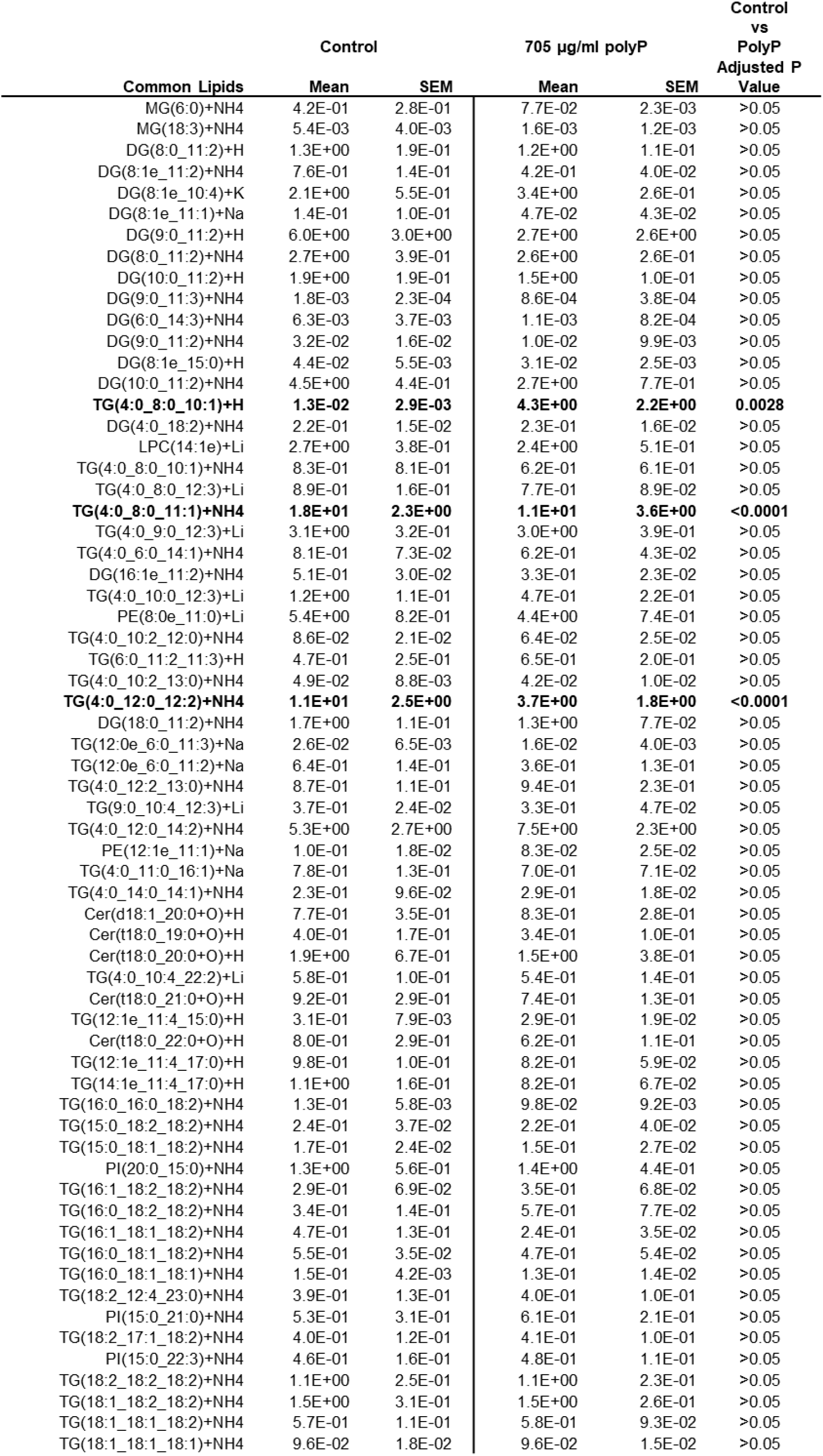
Lipids in Triton X-100 insoluble membranes from WT *D. discoideum* cells in the absence (Control) or in the presence of 705 μg/ml polyP. Relative abundance of common lipids in control and polyP treated Triton X-100 insoluble membranes are shown. Values are mean and SEM from 3 independent experiments. P values from 2 way ANOVA comparison of mean relative abundance from control and polyP treated Triton X-100 insoluble membranes are shown. The lipids that are significantly different are highlighted in bold. MG indicates monoglycerides, DG diglycerides, TG triglycerides, PE phosphatidyl ethanolamine, Cer Ceramides, and PI phosphatidylinositols.

**Table 2:**
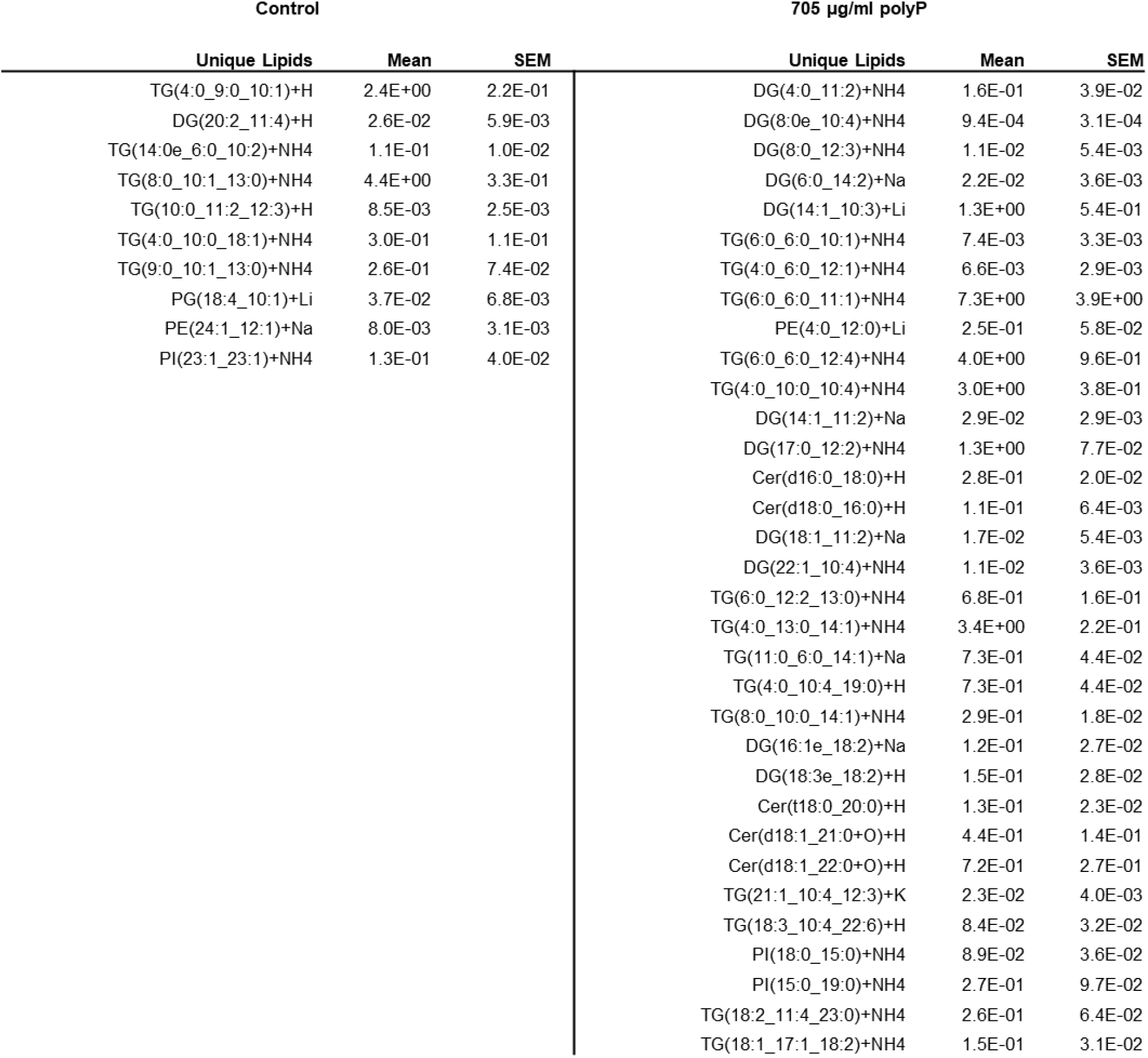
Lipids in Triton X-100 insoluble membranes from WT *D. discoideum* cells in that were detectable only in the absence (Control) or in the presence of 705 μg/ml polyP. The relative abundance of unique lipids in control or polyP treated Triton X-100 insoluble membranes are shown. Values are mean and SEM from 3 independent experiments.

### PolyP alters the protein composition of lipid microdomains

The biophysical and biochemical microenvironment of the membrane microdomains, such as membrane fluidity and lipid composition, affects the protein composition in the microdomains and the downstream signal transduction pathways triggered by microdomain proteins (Lucero and Robbins, 2004). To determine if polyP alters microdomain proteins in addition to fatty acids and lipids, WT cells were incubated with or without 705 μg/ml polyP for 30 minutes and lysed with 1% Triton X-100. The Triton-insoluble extracts were then assayed by proteomics. PolyP significantly increased the accumulation of 80 proteins that fall under the group possessing oxidoreductase activity and 356 proteins under the catalytic activity group based on gene ontology by molecular function (Supplementary Table 1). PolyP significantly increased 344 membrane proteins and 666 cellular anatomical entity proteins, and significantly decreased 637 intracellular anatomical structural proteins in the samples (Supplementary Table 1). Among the detected proteins, polyP significantly upregulated at least 3 known lipid raft proteins and reduced at least 4 lipid raft proteins (Table 3) (Barylko et al., 2009; Eckert and Muller, 2009; Funatsu et al., 2000; Otto and Nichols, 2011; Rauchenberger et al., 1997; Wang and Schey, 2015; Wienke et al., 2006). Together, these data suggest that, in addition to fatty acids and lipids, polyP alters the composition of lipid raft proteins in the membrane microdomains.

**Table 3:** Polyphosphate-induced changes in known lipid raft proteins in Triton X-100 insoluble material. WT *D. discoideum* cells were cultured with or without 705 μg/ml polyP for 30 minutes. The Triton X-100 insoluble material was then analyzed by proteomics. Known lipid raft proteins that are upregulated or downregulated by polyP treatment are listed.

| Upregulated lipid raft proteins |  |  | Downregulated lipid raft proteins |  |  |
| --- | --- | --- | --- | --- | --- |
| Accession ID | Proteins | P-value | Accession ID | Proteins | P-value |
| P54677 | Phosphatidylinositol 4-kinase | <0.05 | Q54WZ2 | Vacuolin-B | <0.05 |
| Q54DL7 | von Willebrand factor A domain-containing protein | <0.05 | O96923 | Gelsolin-related protein of 125 kDa | <0.05 |
| P36412 | Ras-related protein Rab-11A | <0.05 | Q9GPM4 | Phosphoglycerate kinase | <0.05 |
|  |  |  | Q54ET2 | Presenilin-A | <0.05 |

## Discussion

Proliferating *D. discoideum* cells accumulate extracellular polyP when their local cell density increases, and a high concentration of extracellular polyP inhibits proliferation when cells are about to overgrow their food supply (Suess and Gomer, 2016; Suess et al., 2019). Here we found that the extracellular polyP strongly inhibits exocytosis of undigested or partially digested food, but only slightly inhibits the ingestion of food in *D. discoideum*. We envision that this response to extracellular polyP would allow proliferating *D. discoideum* cells to store food rather than digest it when cells are about to starve. A different *D. discoideum* secreted factor called autocrine proliferation repressor AprA also inhibits *D. discoideum* proliferation at high cell densities but does not significantly affect the growth of the cells (Brock and Gomer, 2005). This suggests that the cells find it important to inhibit proliferation but not growth at high cell densities, and use two different factors to accomplish this. Whether AprA inhibits exocytosis is however unknown. Although polyP reduces cell membrane fluidity and alters the composition of lipids and proteins in the lipid microdomains, how polyP changes the membrane physical properties, and whether this directly inhibits exocytosis, is unclear.

We found that polyP requires GrlD, PPK1, and I6kA to reduce macropinocytosis and exocytosis. Possibly because macropinocytic and exocytic activities require fluidic membrane (Degreif et al., 2019), polyP requires GrlD, PPK1, and I6kA to reduce membrane fluidity, indicating that polyP uses a signal tranduction pathway to alter membrane physicalproperties and membrane trafficking. Although cells lacking GrlD and PPK1 display growth and developmental defects (Livermore et al., 2016; Suess and Gomer, 2016), cells lacking I6kA proliferate faster and show normal development (Desfougeres et al., 2022). How *D. discoideum* cells separately use these proteins to rapidly response to polyP is intriguing, and this requires further investigation.

*D. discoideum* cells maintain their cell surface area by coordinating the internalization of the cell membrane and exocytosis of the membrane precursor vesicles (Thilo and Vogel, 1980). Although polyP slightly reduced macropinocytosis, where the internalization of the cell membrane with food particles occurs, polyP inhibited exocytosis to a greater extent. Since exocytosis involves the removal of undigested food particles to the extracellular space and the delivery of precursor membranes to the cell surface, which involves the translocation of exocytic vesicle marker p80 to the cell surface (Charette and Cosson, 2006), inhibiting exocytosis can cause the cells to accumulate internalized membranes and reduce cell surface translocation of p80. *D. discoideum* cells lacking protein kinase C (*pkcA^−^*) accumulate more extracellular polyP and exhibit reduced pinocytosis, but increased exocytosis (Umachandran et al., 2022), suggesting that pinocytosis and exocytosis are not coupled, and polyP might require PkcA to inhibit exocytosis. Continuous inhibition of exocytosis would shrink the cell, but we previously observed that polyP inhibits cytokinesis to increase the percentage of large cells (Suess and Gomer, 2016). How *D. discoideum* maintains cell size while inhibiting exocytosis and membrane recycling is unclear.

At low cell densities, and thus low extracellular polyP concentrations (~15 μg/ml), approximately 0.1 percent of *D. discoideum* cells stop the killing of ingested *E. coli* without affecting the ingestion of the bacteria (Rijal et al., 2020). At high cell densities, when the polyP concentration reaches ~705 μg/ml, ~1.7 percent of *D. discoideum* cells stop the killing of ingested *E. coli*, and many *D. discoideum* cells stop ingesting zymosan bioparticles and stop exocytosing undigested or partially digested zymosan bioparticles to an even greater extent, suggesting a dose-dependent effect of polyP on the storage of food during starvation. Low concentrations of polyP (500 nM) increase mTOR activity (Wang et al., 2003), and activated mTOR inhibits autophagy, perhaps as a mechanism to inhibit the killing of ingested bacteria.

Some lipids and proteins can cluster into discrete microdomains in the cell membrane, called lipid rafts (Levental et al., 2020), and these microdomains are involved in physiological processes such as membrane trafficking (Lingwood and Simons, 2010; Lorent et al., 2017; Simons and Ikonen, 1997; Tooze et al., 2001). Our results indicate that polyP alters the lipid and protein composition of Triton X-100-insoluble lipid microdomains, which may alter membrane fluidity by altering the kinetic stability of the membrane (Grimm et al., 2006), and membrane recycling by altering endocytosis and exocytosis (Ben-Dov and Korenstein, 2013; Ge et al., 2010). However, lipid microdomains are also present in non-plasma membrane systems such as ER, Golgi, and mitochondria (Browman et al., 2006; Garofalo et al., 2005; Wang et al., 2020), suggesting that, in addition to the plasma membrane, polyP might alter homeostasis of sub cellular organelles. Our results do not, however, exclude the possibility that polyP might directly interact with other membrane components, beside lipid microdomains, to alter cell membrane fluidity and membrane recycling.

Lipid microdomains are the hub for various signaling pathways in living cells (Nichols, 2003; Roh et al., 2014). Changes in lipid composition of lipid microdomains has been observed in immune cells, such as T cells, treated with docosahexaenoic acid (Li et al., 2005) and glucocorticoid steroid hormones (Van Laethem et al., 2003). Similar to glucocorticoids, which inhibit endocytosis in a macrophage-like cell line (Miller and Melnykovych, 1982), the high levels of extracellular polyP that allow cells sense that they are at a high cell density and about to overgrow their food supply and starve, might alter lipids and proteins in lipid microdomains to inhibit exocytosis and thus store ingested nutrients in anticipation of starvation.

## Acknowledgements

The use of the Microscopy and Imaging Center facility at Texas A&M University is acknowledged. The Olympus FV1000 confocal microscope acquisition was supported by the Office of the Vice President for Research at Texas A&M University. Use of the TAMU/ Laboratory for Biological Mass Spectrometry service and collaboration facility (LBMS) is acknowledged. This work was supported by National Institutes of Health grants GM118355 and GM139486.

## Author Contributions

R. R. designed, performed experiments, analyzed data, and wrote the paper. S. A. K. and R. J. R. performed experiments, and R. H. G. coordinated the study and wrote the paper.

## Contact for Reagent and Resource Sharing

Further information and requests for reagents may be directed to, and will be fulfilled by, the authors Ramesh Rijal and Richard Gomer.

## Materials and Methods

### D. discoideum cell culture

WT AX2 (DBS0237699) (Fey et al., 2013), *grlD*^−^ (DBS0350227) (Tang et al., 2018), *ppk1^−^* (DBS0350686) (Livermore et al., 2016), *i6kA^−^* (DBS0236426) (Luo et al., 2003), and *i6kA^−^/i6kA* (Suess and Gomer, 2016) *D. discoideum* strains were obtained from the *Dictyostelium* Stock Center. Cells were grown at 21 °C in a type 353003 tissue culture dish (Corning, Durham, NC) in SIH defined minimal medium (Formedium, Norfolk, England) or on SM/5 agar (2 g glucose, 2 g bactopeptone (BD, Sparks, MD), 0.2 g yeast extract (Hardy Diagnostics, Santa Maria, CA), 0.2 g MgCl_2_.7H_2_0, 1.9 g KH_2_PO_4_, 1 g K_2_HPO_4_ and 15 g agar per liter) (http://www.dictybase.org/) on lawns of *E. coli* DB (DBS0350636) in a type 25384-302 petri dish (VWR, Radnor, PA). 100 μg/ml dihydrostreptomycin and 100 μg/ml ampicillin were used to kill *E. coli* in *D. discoideum* cultures obtained from SM/5 agar (Brock and Gomer, 1999). *grlD*^−^, *ppk1*^−^, and *i6kA*^−^ cells were grown under selection with 5 μg/ml blasticidin, and *i6kA^−^/i6kA* cells were grown under selection with 5 μg/ml G418. *D. discoideum* cells from a 80% - 90% confluent culture in a tissue culture dish were collected using a glass pipette, transferred to 15 ml conical tubes (Falcon, VWR), washed 2 times with SIH by centrifugation at 500 x g for 5 minutes, the cell density was measured with a hemocytometer, and 100 μl of cells at 10^6^ cells/ml was transferred to a type 353219 96-well, black/clear, tissue culture treated plate (Corning) to obtain 10^5^ cells per well, or 1 ml was transferred to type 353047 24-well tissue culture plate (Corning) to obtain 10^6^ cells per well. For proliferation assay, *D. discoideum* cells were grown in liquid shaking culture and cell densities were determined as previously described (Suess and Gomer, 2016).

### Bacterial culture

*E. coli* K-12 (BW25113) (CGSC#7636) (Baba et al., 2006; Datsenko and Wanner, 2000) were grown at 37 °C in Luria-Bertani (LB) broth (BD, Sparks, MD). *E. coli* DB were grown at 21 °C on SM/5 agar.

### Recombinant polyphosphatase purification and polyP concentration measurement

Recombinant *S. cerevisiae* exopolyphosphatase (PPX) (Gray et al., 2014; Wurst and Kornberg, 1994) was purified as previously described (Brock and Gomer, 2005). PPX was used for treatment of *D. dicoideum* cells and culture supernatants as previously described (Rijal et al., 2020). Extracellular polyP secreted by *D. discoideum* strains was assessed by adding 25 μg/ml of DAPI (Biolegend, San Diego, CA), and measuring fluorescence at 415/550 nm (excitation/emission) as previously described (Aschar-Sobbi et al., 2008). Culture supernatants were clarified by centrifugation at 12,000 × g for 2 minutes. PolyP concentrations were determined using polyP standards (Spectrum, Cat# S0169, New Brunswick, NJ), and used for all the assays. PolyP (Spectrum) stocks was prepared in PBM, filter sterilized and used for all assays, and was labeled polyP unless specified in the text. To investigate if any potential small molecule contaminants present in the polyP affect an assay, the Spectrum polyP was desalted using a Vivacon 500 2 kDa cutoff spin filter (Sartorius, Bohemia, NY) at 7500 x g for 60 minutes at room temperature, the retentate was resuspended in PBM and collected in an eppendorf tube, and was labeled 2-kDa filtered polyP. Fractionated polyP (medium chain polyP p100 and short chain polyP) were from Kerafast (Boston, MA). PolyP standards of specific chain lengths (60-mer polyP) were kindly provided by Dr. Toshikazu Shiba (RegeneTiss Inc., Japan).

### Resolution of polyP by PAGE and toluidine blue staining of polyP in gel

PolyP was resolved by polyacrylamide gel electrophoresis (PAGE) using a 5.5 × 7.5 cm^2^ 10 % polyacrylamide (Acryl/Bis 19:1 40 % (w/v) solution, VWR Life Science Seradigm, Randor, PA) gel as previously described (Losito et al., 2009; Smith and Morrissey, 2007). The running buffer was 1X TAE (4.84 g Tris, 1.14 ml glacial acetic acid, and 0.37 g Ethylenediaminetetraacetic acid (EDTA) per liter) (all reagents were from VWR Life Science Seradigm, Randor, PA), and the 6X sample buffer was 0.01% Orange G (Fisher Scientific, Fair Lawn, NJ); 30 % glycerol; 10 mM Tris pH 7.4; and 1 mM EDTA. PAGE was performed at 100 V for 1 hour at room temperature until the orange G had run through two thirds of the gel. Gels were stained with 0.05% toluidine blue (Fisher Scientific, Fair lawn, NJ), 20% methanol (VWR) and 2% glycerol for 1 hour, destained 3-4 days with several changes of destaining solution (staining solution without toluidine blue), and images were taken in white light using a Biorad scanner (Bio-Rad, Hercules, CA).

### Endocytosis and exocytosis assays

Endocytosis and exocytosis of TRITC-dextran (average molecular weight 65,000-85,000) (Sigma, St Louis, MO) were performed as previously described (Rivero and Maniak, 2006), with the following modifications. *D. discoideum* cells were seeded in a 96-well, black/clear, tissue-culture-treated plate. After 30 minutes, polyP from a 100 mg/ml stock in PBM was added to the cells and mixed by gentle pipetting. Five microliters of 50 mg/ml TRITC-dextran in SIH was added to a well containing 10^5^ cells and mixed by gentle pipetting, the plates were spun down at 300 × g for 2 minutes in a Heraeus multifuge X3R centrifuge (ThermoFisher Scientific, Germany), and incubated for 30 minutes to allow endocytosis of TRITC-dextran. Cells were washed two times by gently removing the medium, and gently adding 200 μl SIH, fixed with 4% paraformaldehyde (PFA) in PBS for 10 minutes, washed twice with 200 μl of phosphate-buffered saline (PBS), and images were taken with a 100× oil-immersion objective on a Nikon Eclipse Ti2 microscope to measure fluorescence of endocytosed TRITC-dextran. For exocytosis, similar to endocytosis, cells were allowed to endocytose TRITC-dextran for 30 minutes in the absence of polyP, uningested TRITC-dextran was removed by washing twice with 200 μl SIH, and cells were incubated in SIH containing different concentrations of polyP to allow exocytosis of ingested TRITC-dextran. After 30 minutes of incubation, cells were washed with 200 μl of SIH, fixed, and images were taken to determine the fluorescence of ingested TRITC-dextran. Similarly, bulk exocytosis of TRITC-dextran in the presence of polyP of different sizes and purity was measured as previously described (Rivero and Maniak, 2006). The apparent exocytosis of TRITC-dextran slowed down after 3 hours possibly due to reingestion of exocytosed dextran. To overcome this, cells were diluted after 3 hours in SIH to 1/10 of the original density, and were allowed to exocytose TRITC-dextran for 24 hours, and fluorescence of the retained TRITC-dextran per cell was measured.

### Bacterial survival assay and phagocytosis

*E. coli* K-12 survival assays were performed as previously described (Rijal et al., 2020), except that polyP was added to the assay after WT *D. discoideum* cells were allowed to ingest *E. coli* for 4 hours, and the numbers of viable ingested *E. coli* at 24 hours and 48 hours were determined. Phagocytosis of alexa fluor 594 conjugated zymosan bioparticles (ThermoFisher, Rockford, IL) was performed as described in (Rijal et al., 2020).

### Immunofluorescence

p80 staining was performed as previosuly described (Charette and Cosson, 2006). For total p80 staining, *D. discoideum* cells were seeded in a 96 well, black/clear, tissue culture treated plate, spun down at 500 × g for 2 minutes, and the SIH media was changed for SIH containing polyP. After 30 minutes of incubation, cells were fixed with 4% PFA for 10 minutes. In a control, at time 0, the media supernatant was discarded and cells were fixed with 4% PFA. Cells were washed three times with 200 μl of PBS, and permeabilized with 0.1 % Triton X-100 (Alfa Aesar, Tewksbury, MA) in PBS for 5 minutes. Cells were washed two times with PBS, blocked with 1 mg/ml type 0332 bovine serum albumin (VWR) in PBS for 1 hour and washed once with PBS. 1:200 anti-p80 antibody (H161) (Developmental studies hyrbidoma bank) in PBS/0.1 % Tween 20 (Fisher Scientific, Pittsburgh, PA) (PBST) was added to cells and incubated at 4 °C overnight. For surface p80 staining, cells were incubated in media containing polyP for 30 minutes, media supernatant was replaced with ice-cold media containing polyP and incubated for 5 minutes, 1 μl of anti-p80 antibody (H161) (Ravanel et al., 2001) was added to the cells and incubated for 10 minutes, cells were washed twice with ice-cold media to remove excess unbound antibodies, and fixed with 4% PFA for 10 minutes. Cells stained for total or surface p80 were washed three times with PBST, and incubated with 1:500 Alexa 488 anti-rabbit (Jackson Immunoresearch, West Grove, PA) in PBST for 1 hour. Cells were washed three times with PBST and 200 μl of PBS was then added to the well. For phalloidin staining, cells incubated with polyP for 30 minutes were fixed, permeabilized, and stained with 1:2000 ifluor 555 phalloidin (Abcam, Cambridge, MA) in PBS in the dark for 30 minutes. Cells were washed three times with PBS before taking images. Each washing step was done for 5 minutes and all steps were performed at room temperature if not indicated otherwise. Images of cells were taken with a 100× oil-immersion objective on a Nikon Eclipse Ti2 (Nikon), and deconvolution of images was done using the Richardson-Lucy algorithm (Laasmaa et al., 2011) in NIS-Elements AR software. Fluorescence intensity of p80 was analyzed by Fiji (ImageJ; NIH).

### Cell membrane recylcing

Cell membrane staining was performed as previously described (Tanaka et al., 2017). To observe the cell membrane, WT *D. discoideum* cells were seeded in a 96 well, black/clear, tissue culture treated plate, and spun down at 500 × g for 2 minutes. Cells in SIH medium were incubated for 30 minutes in the absence or presence of polyP, and SIH was replaced with SIH containing 1.25X CellMask green stain (Invitrogen). Ten minutes after staining, cells were washed twice with SIH medium, incubated with SIH containing the indicated concentration of polyP, and cells were excited by an Argon laser (488 nm) and emission was observed through a 515-530 nm band pass filter. Cells were visualized with a 60× water-immersion objective on a FV1000 confocal microscope (Olympus, Center Valley, PA) equiped with Olympus Fluoview Ver.4.2a software, and time-lapse images were taken for 8 minutes with an interval of 30 s. Fluorescence intensity of the cell or the cell interior was measured using ImageJ software, and corrected total fluorescence was calculated as integrated density – (Area of selected cell x mean fluorescence of background readings). The fluorescence intensity at 0 time was set to 1.

### Fluorescence recovery after photobleaching

Photobleaching was performed as previously described (Tanaka et al., 2017). In brief, *D. discoideum* cells were seeded in a 96 well, black/clear, tissue culture treated plate, and spun down at 500 × g for 2 minutes. Cells in SIH medium were incubated for 30 minutes in the absence or presence of the indicated concentration of polyP, SIH was replaced with SIH containing 1.25X CellMask green stain (Invitrogen). Ten minutes after staining, cells were washed twice with SIH medium, incubated with SIH containing the indicated concentration of polyP, and the full power of the argon laser (488 nm) was applied to a region of a cell for 0.5 second. Images were taken every 0.5 s for 1 minute with a 60× water-immersion objective on a FV1000 confocal microscope (Olympus, Center Valley, PA) equiped with Olympus Fluoview Ver.4.2a software. A circular region of interest (ROI) of 3 μm diameter in the cell membrane was bleached, and the fluorescence intensity in the photobleached area (FRAP ROI) was monitored over time. The fluorescence intensity in a 3 μm diameter unbleached region outside of the cell (Base ROI) and a control cell (reference cell ROI) were also monitored. 10 images (10 frames in 5 seconds) were taken before the FRAP ROI was photobleached to get pre-bleached fluorescence intensities for all ROIs. Normalized fluorescence recovery, half-life of recovery, diffusion coefficient, and mobile fraction were then calculated following (Eils and Kappel, 2004; Tanaka et al., 2017).

### Scanning electron microscopy

For scanning electron microscopy, cells were seeded in 3 ml SIH medium on a glass coverslip on the bottom of a type 353046 6-well plate (Corning) for 30 minutes, 705 μg/ml polyP or 100% CM was added to cells and incubated for 30 minutes, and the medium was gently transferred to an eppendorf tube containing 120 μl 25% glutaraldehyde (Sigma), mixed, gently added back to the cells, and incubated for 30 minutes at room temperature. The final concentration of glutaraldehyde was 1%. Cells were rinsed with PBS and dehydrated by successive 30 minutes incubations with 30%, 50%, 70%, 90%, 100%, 100%, and 100% ethanol with a very gentle agitation on a orbital shaker. Cells were not allowed to dry during this procedure. The ethanol was replaced with a Ethanol: Hexamethyldisilazane (HMDS) (Sigma) mixture at a 1:1 and then a 1:3 ratio, and then pure HMDS was added. Each incubation step was 2 hours with gentle agitation. All work was done in a fume hood. Pure HMDS was allowed to evaporate overnight to dry the cells. After coating with 20 nm gold using a type 108 sputter coater (Cressington, Redding, CA), the cells were observed with a scanning electron microscope (Tescan Vega, Warrendale, PA).

### Lipid microdomain extraction, fatty acid analysis, and lipid and potein analysis

Detergent-dependent lipid raft isolation was done following (Lingwood and Simons, 2007) with the following modifications. *D. discoideum* cells at 10^6^ cells/ml density were incubated in 10 ml SIH in the absence or presence of 705 μg/ml polyP. After 30 minutes, cells were collected by centrifugation at 500 × g for 5 minutes and resuspended in 1 ml TNE buffer (150 mM NaCl, 2 mM EDTA, 50 mM Tris-HCl, pH 7.4) containing protease and phosphatase inhibitors (Cell Signaling Technology, Danvers, MA), passed through a 25 gauge needle to shear the cells, and then treated with 1% Triton X-100, and after 30 minutes of incubation on ice, lysates were centrifuged at 15,000 × g for 1 hour at 4°C. For lipid extraction to detect fatty acids and lipid contents of the 1% Triton X-100 insoluble membrane fractions, the supernatant was discarded, and the pellet was resuspended in 750 μl of TNE buffer containing protease and phosphatase inhibitors and mixed with an equal volume of 2:1 methanol:chloroform solution. The mixture was vortexed for 10 minutes at room temperature, centrifuged at 1000 × g for 3 minutes, the lower organic layer was transferred to a 5 ml Wheaton glass vial, and the liquid was allowed to dry overnight in a vacuum dryer. The extracts were sent to the Department of Chemistry Mass Spectrometry core facility at Texas A&M University for gas chromatography-mass spectrometry (GC-MS) on a DSQ II (Thermo Scientific, TX), and liquid chromatography-mass spectrometry (LC-MS) on a QE Focus (Thermo Scientific) for fatty acid and lipid analysis, respectively. For fatty acid analysis, lipid extracts were transesterified to methyl esters as described in (Metcalffe and Wang, 1981). Decanoic acid was used as an internal control for fatty acid quantitation and its abundance was normalized to a value of 1. Abundance of identified fatty acids were determined as relative to Decanoic acid. The amount of each lipid was calculated as the percent area of the sum of all the lipids.

For protein content determination, the 1% Triton X-100 insoluble fraction (pellet) was resuspended in 1X SDS sample buffer containing protease and phosphatase inhibitors, and heated at 95 °C for 5 minutes. Samples were loaded onto 4-20% polyacrylamide gel, electrophoresed until the bromophenol blue in the sample buffer had migrated 5 mm from the bottom of the wells, the portion of the gel from the bottom of the well to the dye front was excised with a clean razor blade, diced into 2 mm^2^ cubes, and transferred to Eppendorf tubes pre-rinsed with ethanol. Sample preparation for LC-MS was performed using in-gel protein digestion protocol at the Department of Chemistry mass spectrometry core facility at Texas A&M University (https://mass-spec.chem.tamu.edu/proteomics/proteomics-protocols.php). Mass spectrometry proteomics was performed on a Thermo Scientific Orbitrap Fusion tribrid mass spectrometer equipped with a Dionex UltiMate 3000 reverse-phase nano-UHPLC system. Biological processes, molecular functions and cellular locations were determined for identified proteins using the Gene ontology resource (http://geneontology.org/), and PANTHER analysis and enrichement analysis was performed using Fisher’s Exact test and Bonferroni correction for multiple testing. Membrane raft proteins were identified in Triton X-100 insoluble membrane fraction by searching the Uniprot database and pubmed.

### Cell motility, speed, and directionality measurement

Cells were seeded in 96 well, black/clear, tissue culture treated plates, spun down at 500 × g for 2 minutes, and SIH media was changed to SIH, SIH containing polyP, or 100% CM. At least 30 cells per experiment were imaged every 15 seconds for 30 minutes using a 40× objective on a Nikon Eclipse Ti2 (Nikon). Accumulated distance, speed and directionality was calculated as previously described (Phillips and Gomer, 2012).

### Cytoskeletal protein extraction and immunoblotting

Cells were cultured as above, and after 30 minutes, for whole cell lysates, the culture medium was removed and cells were lysed in 75 μl of 1X SDS sample buffer. Crude cytoskeletons were extracted as previously described (Rijal et al., 2019). The whole cell lysate and crude cytoskeletons were resolved on 4-20% Mini-PROTEAN tris-glycine polyacrylamide gels (Bio-Rad), and Western blots were stained with 1:5000 diluted anti-beta-actin mouse monoclonal antibody (Cell Signaling Technology) to detect total and cytoskeletal actin as described in (Rijal et al., 2019).

## Quantification and Statistical Analysis

### Statistical analysis

Statistical analyses were performed using GraphPad Prism 9 (GraphPad, San Diego, CA). A p < 0.05 was considered significant.

